# UPLiFT: Uncertainty Parameterized Activation for Lightweight Fine-Tuning

**DOI:** 10.64898/2026.10.05.756747

**Authors:** A N M Nafiz Abeer, Amir Hossein Rahmati, Xiaoning Qian, Byung-Jun Yoon

**Affiliations:** Texas A&M University; Brookhaven National Laboratory

## Abstract

Learned protein representations contribute to the broad utility of pretrained protein language models (PLMs), often enhanced by task-specific fine-tuning where parameter-efficient techniques are imperative due to large network size. Our work takes an orthogonal approach to such fine-tuning: rather than optimizing the network weights, we formulate adaptation as an optimization task over the parameterized activation space. Motivated by the prior work on curvature-tuning, we propose **UPLiFT, U**ncertainty **P**arameterized activation for **Li**ghtweight **F**ine-**T**uning, with enhanced adaptation capacity when the underlying model—whether pretrained or randomly initialized—remains fixed, for significantly improved parameter efficiency. Furthermore, its learnable non-linearity naturally allows to model uncertainty over the activation space through fine-tuning in a variational inference framework. Evaluation across multiple protein datasets shows that UP-LiFT performs competitively while requiring 3 − 11 fewer trainable paramters than weight-only adaptations. Bayesian UPLiFT further improves calibration while largely preserving the predictive performance of deterministically fine-tuned models. The narrow predictive performance margin with the low-rank adaptation baseline in residue-level tasks highlights the potential of our proposed approach as a parameter-efficient fine-tuning technique for PLM-based predictors.

## 1 Introduction

The learned knowledge as embedded representations in the protein language models (PLMs) (Lin et al., 2023; Hayes et al., 2025) through self-supervised pretraining with large-scale datasets of protein sequences has led to PLMs’ significant applications covering broad spectrum, ranging from sequence-structure-function relationships to *de novo* design, encompassing protein property prediction and variant effect prediction (Notin et al., 2023; Dinh et al., 2026; Marquet et al., 2022), structure prediction (Lin et al., 2023; Candido et al., 2026), generative molecular design (Tran et al., 2026; Huang et al., 2025), and more (Abdine et al., 2024; Simon & Zou, 2025). While pretrained PLMs offer informative representation space useful for the downstream tasks, fine-tuning their parameters often refines the representations with consistent improvement across diverse downstream tasks (Schmirler et al., 2024; Zhang et al., 2026a; Abeer et al., 2025; Calef et al., 2026; Bushuiev et al., 2026; Dinh et al., 2026).

However, full parameter adaptation increases computational overhead relative to simply training only a task-specific prediction head utilizing fixed PLM embeddings. Consequently, for larger models, one has to opt for parameter-efficient approaches such as Low-Rank Adaptation (LoRA) instead of fine-tuning full models (Hu et al., 2022; Schmirler et al., 2024). Orthogonal to such established approaches from large language models (LLMs), Hu et al. (2026) recently proposed *curvature tuning* (CT) that fine-tunes the parameterized activation function of a pretrained network without altering its weights. Their results indicate that the activation space of a pretrained network has a more parameter-efficient yet controllable mechanism facilitating adaptation to downstream tasks in broadly similar capacity as LoRA. While this CT approach was not applied in the protein domain, a similar trend is expected to hold for PLMs, specifically given the growing field of works on activation steering for targeted protein sequence design (Huang et al., 2025; Simon & Zou, 2025).

Our work builds on this premise of training parameterized activation functions by extending existing approach along two fronts. First, Hu et al. (2026) learned the activation parameters of pretrained network together with the weights of the task-specific prediction head, whose activation response remained fixed. We asked: what if we could improve the learning capacity of the parameterized activation function, moving beyond steering to activation-only adaptation? Consequently, this could lead to increased parameter efficiency as task-specific fine-tuning could be done by leveraging only the activation space of both the pretrained network and prediction head, even though the latter is frozen to randomly initialized weights.

Second, the reliability of PLM-embedding-based predictors can be enhanced through seamless integration of uncertainty quantification (UQ) approach (Greenman et al., 2025; Groth et al., 2024; Qayyum et al., 2026) without the additional learnable parameter overhead common to most of the existing weight-space uncertainty methods (Yang et al., 2024; Wang et al., 2024; Rahmati et al., 2026; Zhang et al., 2026b). Combining these two intertwined directions, we aimed to equip the activation-based fine-tuning method with UQ capability over the activation space of the PLM. Accordingly, we first formulated a parameterized activation function with affine-like behavior that provides increased control of gain and offset compared with CT’s modulation around the origin. By imposing a Beta distribution over the bounded curvature parameter of our proposed activation function, we model uncertainty over the activation space through task-specific fine-tuning.

The activation-only nature of our proposed method leads to improved parameter efficiency while maintaining performance across multiple protein datasets, compared to weight-only adaptation strategies such as LoRA and prediction head-only training. To summarize our work,

- we propose, UPLiFT, a parameterized activation function as a fine-tuning plug-in with improved adaptation capacity over existing curvature tuning approach under fixed network weights;
- we formulate this parameterization to naturally model uncertainty over the activation space during task-specific fine-tuning;
- we evaluate the activation-only fine-tuning approach for a wide range of protein datasets, demonstrating competitive performance, with a more pronounced effect in residue granularity, while requiring substantially fewer parameters than weight-space adaptation baselines.

## Adaptation through activation

### Problem formulation

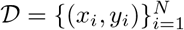 denote a training dataset, and *f*_Θ,*ω*_ denote a model parameterized by its base parameters, Θ, and activation parameters, *ω*. UPLiFT fixes the base model at Θ_0_ and formulates adaptation as optimization over activation space parameters, *ω*,

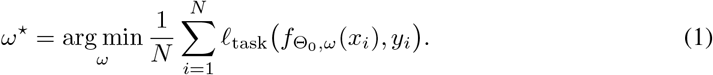

Therefore, the resulting model, 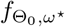, is adapted to the downstream prediction task without any modification to base model parameters, Θ.

### Background

Fine-tuning can change a pre-trained network through two complementary components to improve its prediction: its affine transformations, ***Wx*** + ***b*** with base parameters Θ = {***W***, ***b***}, and its nonlinear activations parameterized by *ω*. Most parameter-efficient methods modify the former (Hu et al., 2022; Lian et al., 2022; Yang et al., 2024). Curvature tuning (Hu et al., 2026) explores the latter, but its activation unit has limited capacity to compensate for the gap originating from *all* affine parameters being frozen.

For any input *z*, curvature-tuning (CT) parameterizes the activation function as

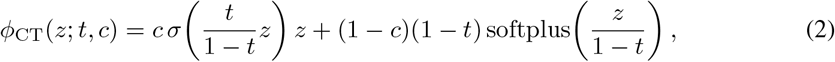

where the parameters *t, c* ϵ (0, 1) can control the activation around its transition regions. However, its positive tail converges to the identity mapping, *ϕ*_CT_(*z*; *t, c*) = *z* + *o*(1) as *z* →+ ∞. Consequently, when the base model parameters (Θ) are fixed, this formulation cannot independently change the asymptotic gain or introduce an offset. To showcase this, we construct a controlled experiment where a small multilayer perceptron (MLP) is randomly initialized, after which every weight and bias is frozen. As demonstrated in Figure 1, CT performs well on the linearly separable task; however, it is less effective on the nonlinear two-moon task demonstrating its limitations.

**Figure 1:**
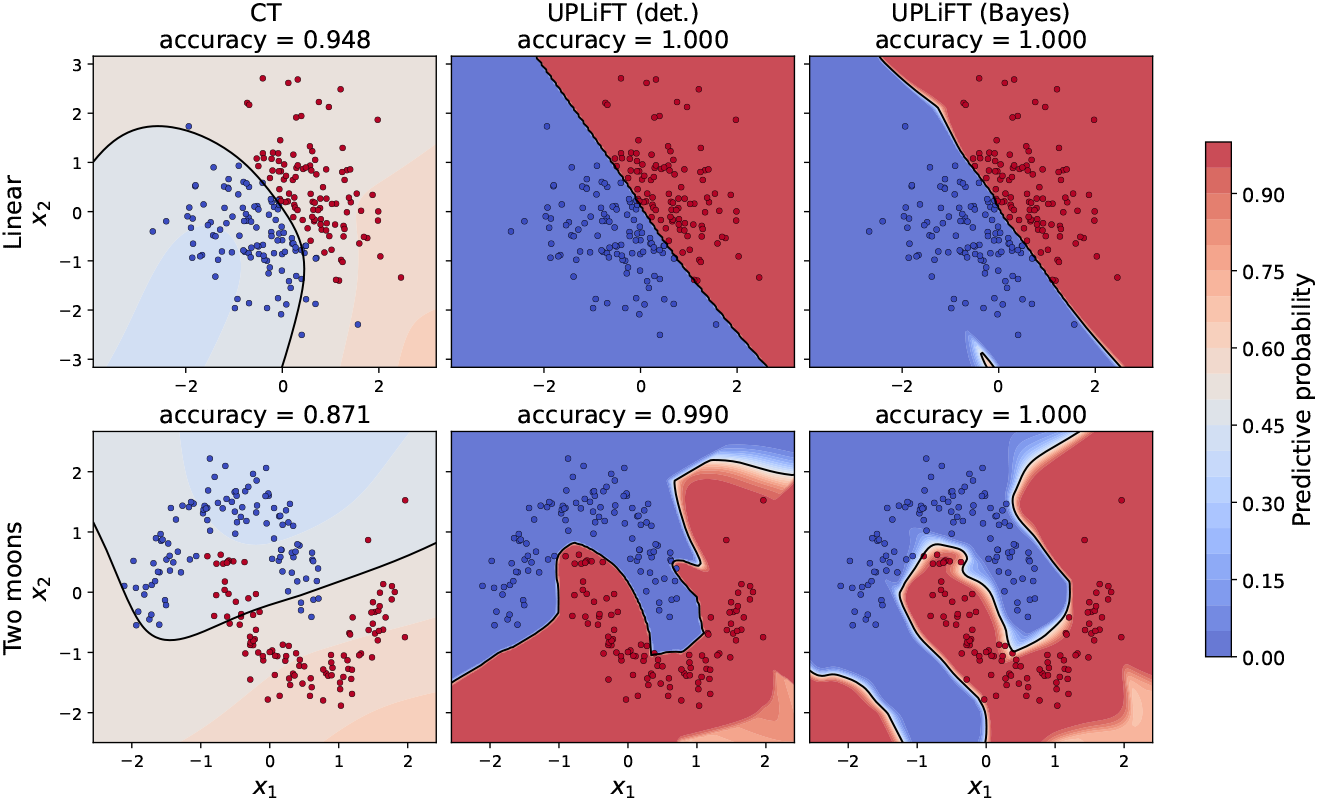
Activation-only adaptation. The same randomly initialized MLP is used for every method, and all weights and biases remain frozen while only activation parameters are optimized. **Top** and **bottom** panels show predictive probabilities and decision boundaries for a linear task and a nonlinear (two-moons) task, respectively. The test samples are shown as scattered points.

Motivated by this observation, we propose **UPLiFT**, which reformulates CT from a predominantly multiplicative transformation, *h*(*z*)*z*, into an additive activation with affine-like behavior. While the positive tail of CT is fixed to *z*, UPLiFT can behave asymptotically as a rectified affine map as detailed later in Section 2.1. This enables activation tuning to control not only where the representation bends, but how it is scaled and shifted across layers by allowing flexible adaptation of the gain and offset at each neuron while leaving the model weights unchanged. UPLiFT aims to make activation-only fine-tuning sufficiently expressive for frozen models. In Section 2.2, we further extend UPLiFT with a principled Bayesian mechanism for uncertainty-aware fine-tuning by probabilistically formulating the bounded curvature parameter. When applying both deterministic and Bayesian UPLiFT variants to the same experiment, trained by updating only their activation parameters as in CT, both UPLiFT variants recover substantially more refined boundaries as shown in Figure 1, consistent with the added gain and offset control provided by UPLiFT.

### 2.1 UPLIFT

Based on the preceding observations, we hypothesize that the limited adaptation flexibility of CT stems from the predominantly multiplicative nature of its transformation, and that an affine-like formulation can provide greater flexibility. We therefore introduce UPLiFT in detail as a new activation-space parameterization designed to increase adaptation capacity.

#### Definition 2.1

(UPLiFT). *For t* ∈ (0, 1) *and c >* 0, *define*

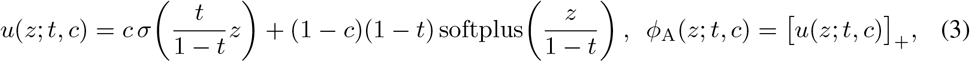

*where* [*v*]_+_ = max(0, *v*). ***UPLiFT*** *applies ϕ*_A_ *independently to every selected hidden feature. The parameters are represented as*

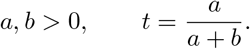

Unlike Equation (2), the bounded sigmoid component in Equation (3) is additive rather than multiplied by *z*. This decouples the transition geometry from the magnitude of the input.

#### Lemma 2.2

(Affine-like tail behavior). *For fixed t* ∈ (0, 1) *and finite c >* 0,

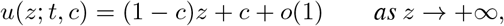

*and*

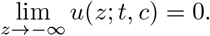

*Therefore*,

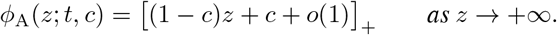

*Proof*. As *z* → +∞, *σ*(*tz/*(1 − *t*)) = 1 + *o*(1) and

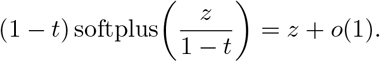

Substitution into Equation (3) gives the first result. Both the sigmoid and softplus terms converge to zero as *z* → −∞, giving the second result. □

Thus, UPLiFT provides a rectified affine-like response [*wz* + *b*]_+_ with effective gain *w* = 1 − *c* and offset *b* = *c*. These quantities are not independent, so UPLiFT is not equivalent to a fully trainable affine layer. Nevertheless, it restores gain- and offset-like control that is absent from the positive tail of CT. Since *c* is not restricted to (0, 1), values *c >* 1 enable extrapolation beyond the convex interpolations explored by CT.

Furthermore, Definition J.1 in the Appendix J shows that, for fixed activation parameters, UPLiFT is globally Lipschitz in its input and can retain nonzero local activation curvature.

#### Representing activation response of pretrained model

The pretrained models often have activation functions with non-zero response for negative inputs, such as GELU. The CT formulation in Hu et al. (2026) allows approximation of GELU through the convex combination of SiLU and Softplus by setting *t* = 0.64 and *c* = 1. To preserve a similar localized negative response around the origin in our parameterized activation function, we introduce a (non-trainable) correction term in 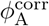 which closely approximates the GELU with *t*=*c*=0, *d*_0_=3.66 and *τ*_0_=0.762 (Appendix K).

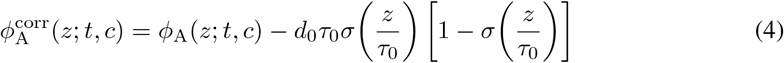

Note that this correction term is localized around the origin and vanishes in both tails. It therefore introduces a GELU-like negative notch without changing the affine-like asymptotic behavior in Definition 2.2, while requiring no additional trainable parameters.

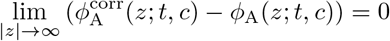

This correction term is important, specially for optimizing over the activation space of a pretrained model. One naturally expects the initialization of activation parameters to closely resemble the activation function that was kept fixed during pretraining since it allows selective adaptation of the activation space guided by the task-specific fine-tuning. Specifically, alignment with the pretrained activation responses when starting the optimization of activation parameters ***ω*** enables effective utilization of the pretrained weights, which can be advantageous for tasks requiring minimal adaptation of pretrained representation. The ablation experiment in Appendix F shows empirical evidence.

### 2.2 Uncertainty Through Stochastic Activations

Rather than deploying a fully Bayesian implementation, UPLiFT quantifies the uncertainty only over the bounded curvature variable *t*, motivated by the effectiveness of activation adaptation. This produces a lightweight activation-space uncertainty model as detailed below.

#### Mean-field activation distribution

For activation feature *j* at layer *l*, let *q*_*l*j_(*t*) = Beta (*t*; *a*_*l*j_, *b*_*l*j_), We use the factorized distribution

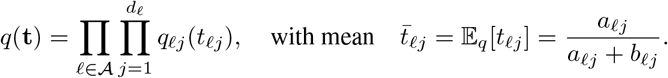

The deterministic UPLiFT evaluates every activation at 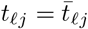. Here, denotes the set of adapted layers and *d*_*l*_ denotes the number of activation features in layer *l*. The distribution *q*(**t**) defines the learned activation uncertainty and its samples are obtained through the following mean-preserving variance control.

##### Definition 2.3

(Variance-controlled sampling). *Let κ* ≥1 *be a concentration multiplier. For each sample, we draw*

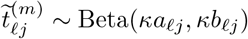

*and define*

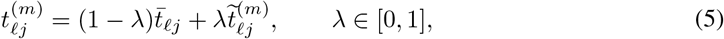

A single vector of activation parameters 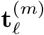 is sampled for each layer using Equation (5) and shared across the inputs in a forward pass. Direct sampling from *q*(**t**) is recovered when *κ* = *λ* = 1; increasing *κ* or decreasing *λ* reduces stochasticity without changing the mean.

##### Lemma 2.4

(Mean preservation and variance control). *After variance-controlled sampling*,

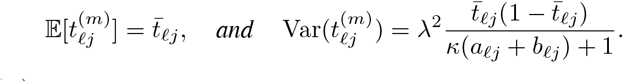

*Consequently*,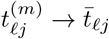 *in mean square as λ* → 0 *or κ*(*a*_*l*j_ + *b*_*l*j_) → ∞.

*Proof*. Scaling both Beta shape parameters by *κ* preserves their ratio and therefore preserves the mean. The variance of Beta(*κa, κb*) is

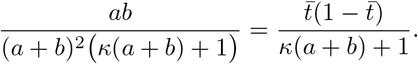

Interpolation with a constant mean multiplies this variance by *λ*^2^. □

#### KL-regularized stochastic objective

Let

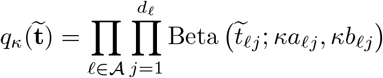

denote the distribution underlying variance-controlled sampling in Definition 2.3. Using prior

Beta(*t*; *a*_0_, *b*_0_), for a minibatch ℬ, then the Bayesian UPLiFT objective is

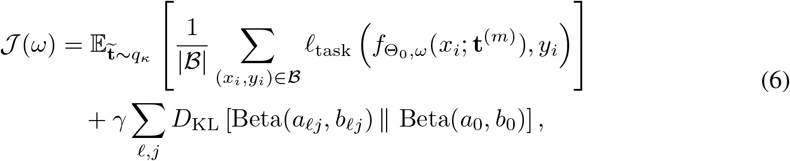

where *γ >* 0 is the regularizing coefficient. The expectation is estimated by Monte Carlo using reparameterized Beta samples.

The expected task loss in Equation (6) favors activation distributions that support accurate prediction, while the KL term regularizes the underlying Beta parameterization toward the prior. Variance control modifies the stochastic activations entering the task loss to promote stable training and limit performance degradation relative to deterministic adaptation. When *κ* = *λ* = 1, samples are drawn directly from *q*(**t**), and the sampling and regularization terms recover the usual mean-field KL-regularized form of variational inference.

#### Posterior predictive approximation

At inference, we draw *M* independent activation samples and perform *M* forward passes while keeping all non-stochastic parameters fixed. For multilabel classification, let 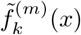 denote the sampled logit for label *k*. The predictive probability is approximated by

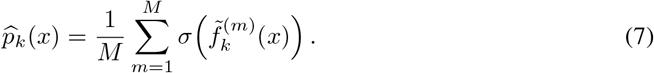

## 3 Results

### 3.1 Experimental settings

#### Prediction tasks

To evaluate the PLM fine-tuning capability of **UPLiFT**, we benchmark multiple tasks involving both local (residue-level) and global (sequence-level) property predictions. Table 1 lists these prediction tasks, among which we have utilized datasets from original works for post-translational modification (PTM) (Zhang et al., 2026a) and normal mode vector (NMV) (Ni & Buehler, 2026). The remaining datasets are collected from the processed version shared in Calef et al. (2026). Appendix B provides more dataset-specific details, such as sequence-similarity restricted data partitioning for NMV.

**Table 1:** Overview of residue-level and sequence-level protein prediction tasks.

| Granularity | Task | Type | Source |
| --- | --- | --- | --- |
| Residue | Post-translational modification (PTM) | Multi-label classification | Zhang et al. (2026a) |
|  | Binding residue categorization | Multi-label classification | Dallago et al. (2021) |
|  | Functional (binding & catalytic) site prediction | Binary classification | Zhang et al. (2024) |
|  | Normal mode vector (NMV) prediction | Regression | Ni & Buehler (2026) |
| Sequence | Binary localization | Binary classification | Almagro Armenteros et al. (2017) |
|  | Subcellular localization | Multi-class classification | Almagro Armenteros et al. (2017) |
|  | Thermostability | Regression | Dallago et al. (2021) |

#### Prediction architecture and adaptation baselines

We evaluate all adaptation mechanisms using the same backbone and prediction architecture. We use the 150M-parameter ESM-2 model (Lin et al., 2023) as the PLM backbone. Residue-level tasks operate directly on the final residue representations, whereas sequence-level tasks use masked mean pooling. The resulting features are processed by a residual prediction head with a 512-dimensional hidden representation and a task-specific output projection. Apart from pooling and output dimensionality, the architecture is fixed across tasks and adaptation methods. See Appendix L for further details on hyperparameters.

We compare deterministic **UPLiFT** with two weight-space baselines. 1) **LoRA tuning** applies rank-16 LoRA adapters (Hu et al., 2022) to the query, key, and value projections of ESM-2 while training the prediction-head weights. 2) **Head tuning** freezes ESM-2 and trains the prediction head. In contrast, **UPLiFT** freezes weights of both the pretrained PLM and the randomly initialized prediction-head, and optimizes only the activation parameters in both. We separately compare **UPLiFT** with the **CT** (Hu et al., 2026) under the same activation-only setting in Section 3.3 and Appendix E.

#### Evaluation protocol and metrics

We select checkpoints according to the task-specific validation criterion and report test performance with the mean and standard deviation over three random seeds. For classification, we use accuracy, F1, and MCC to evaluate discrete predictions, and AUROC and AUPRC to measure threshold-independent discrimination. For regression, Pearson correlations measure predictive association, while MAE, and RMSE quantify prediction error. Probabilistic predictions are also assessed using NLL, ECE, CRPS, and the Brier score in our uncertainty quantification experiments. More detailed definitions and evaluation procedures are provided in Appendix C.

### 3.2 Evaluation of adaptation

#### Activation-space fine-tuning

Table 2 summarizes performance comparison results. Our **UPLiFT** consistently outperformed the **Head tuning** baseline for all residue-level tasks, with demonstrating **LoRA tuning** comparable performance for PTM and NMV. This can be explained by the prominent performance gaps between **LoRA tuning** and **Head tuning**, which indicates the relative importance of adapting the PLM representations over the prediction head. Specifically, the PTM dataset illustrates the most challenging scenario, quantified by the poor performance with the pre-trained PLM. Remarkably, **UPLiFT** demonstrates comparable PTM classification performance to the LoRA fine-tuned models despite training only 231,936 parameters. Given that **UPLiFT** adapts the activation parameters for frozen weights alongside a randomly initialized prediction head, its competitive performance against LoRA on residue-level tasks further demonstrates its capacity to tune PLM’s activation space.

**Table 2:** Adaptation results. Results are reported as mean ±_std_ over three seeds. ↑ / ↓ indicates higher/lower is better. The column for **UPLiFT** is colored according to mean-based performance rank: 1, 2, and 3. Comparisons across more performance metrics are given in Appendix D.

| Task | Metric | LoRA tuning | Head tuning | UPLiFT (ours) |
| --- | --- | --- | --- | --- |
| *Residue-level tasks* |  |  |  |  |
| PTM | F1 ↑ | 0.4089±0.0037 | 0.3124±0.0020 | 0.4080±0.0026 |
| Binding residue categorization | F1 ↑ | 0.3327±0.0061 | 0.3116±0.0067 | 0.3158±0.0074 |
| Binding site binary | AUROC ↑ | 0.8990±0.0011 | 0.8630±0.0016 | 0.8772±0.0013 |
| Catalytic site binary | AUROC ↑ | 0.9371±0.0012 | 0.9010±0.0047 | 0.9194±0.0014 |
| Normal mode vector | Pearson ↑ | 0.6720±0.0053 | 0.6353±0.0031 | 0.6738±0.0035 |
| *Sequence-level tasks* |  |  |  |  |
| Binary localization | Accuracy ↑ | 0.9365±0.0004 | 0.9280±0.0017 | 0.9224±0.0052 |
| Subcellular localization | Accuracy ↑ | 0.8148±0.0025 | 0.7901±0.0078 | 0.7833±0.0051 |
| Thermostability | Pearson ↑ | 0.6852±0.0074 | 0.6894±0.0063 | 0.6586±0.0078 |
| *Parameter efficiency* |  |  |  |  |
| Trainable parameters (range) |  | 2.501–2.506M | 0.657–0.662M | 0.232M |
| Fraction of LoRA tuning |  | 100% | 26.3% | 9.3% |

For sequence-level classification tasks, this proposed activation-only approach was reasonably competitive with the weight-only baselines. Furthermore, **UPLiFT** shows a larger gap for the Thermostability regression task unlike the rest of the datasets. However, this has little practical implication since high predictive performance with the pretrained model made adaptation within PLM unnecessary for this dataset. Particularly, the high performance of **Head tuning** in sequence-level tasks indicates the low sensitivity to residue-level adaptation for learning such global properties. Indeed, **UPLiFT** becomes competitive with **LoRA tuning** (Tables 11 and 12; Appendix G) when we optimize the prediction head weights along with the activation space of PLM.

Figure 2, further illustrates the main trends presented in Table 2 using the mean performance of LoRA-tuned models as the reference (100%) for each task.

**Figure 2:**
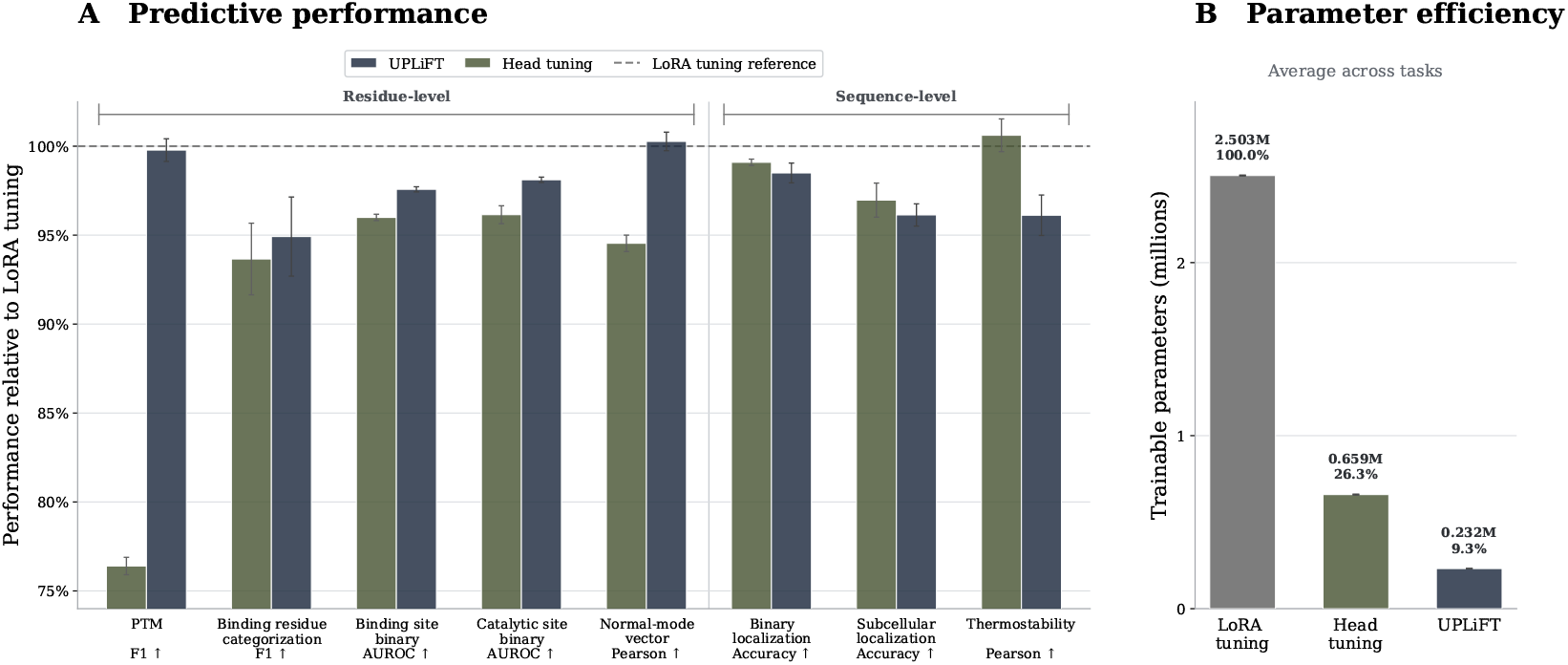
Predictive performance relative to **LoRA tuning** (A), and average trainable parameter counts (B). **UPLiFT** provides competitive performance while using only 9.3% of the trainable parameters compared to **LoRA tuning**.

#### Activation space uncertainty

We evaluate whether the stochastic activation parameterization improves predictive uncertainty while preserving prediction performance. Focusing on residue-level tasks, we compare deterministic and Bayesian **UPLiFT** using task-specific predictive metrics, supplemented by calibration metrics. As shown in Table 3, Bayesian **UPLiFT** achieves lower mean NLL and ECE across all classification tasks. Specifically, Bayesian **UPLiFT** improves mean NLL from 0.2309 to 0.2240 in Binding residue task and mean ECE from 0.0143 to 0.0127 in Catalytic site task, offering better calibrated predictions. The Brier score follows the same trend on three of the four tasks, with the largest improvement in the binding site prediction. Predictive performance is largely retained with Bayesian **UPLiFT**. PTM is particularly notable where Bayesian **UPLiFT** enhances the mean predictive performance (F1 score) from 0.4080 to 0.4107.

**Table 3:**
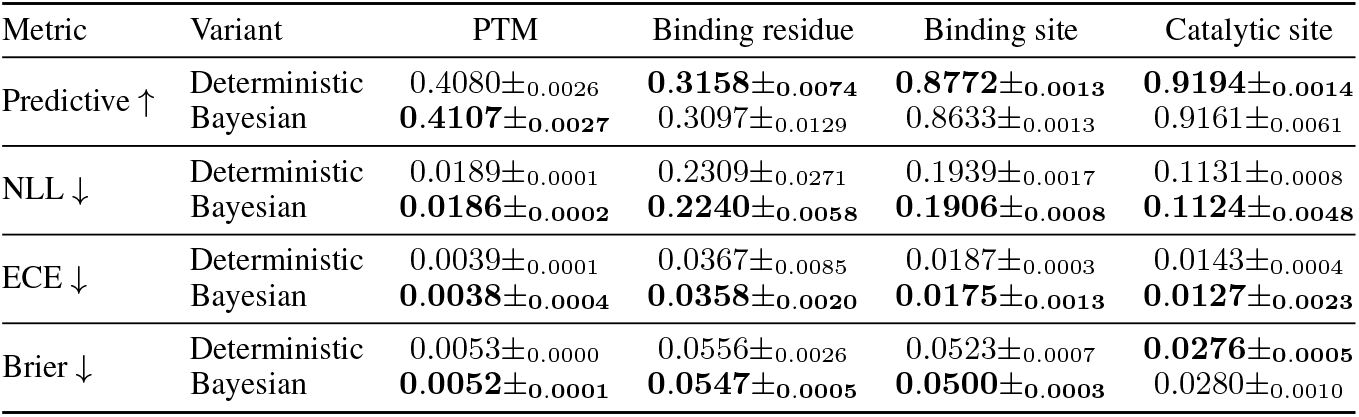
Built-in uncertainty quantification. Performance on uncertainty quantification of Bayesian (*M* = 10 samples) vs. deterministic **UPLiFT**. Reported as mean ±std over three seeds. The best mean performance metrics are highlighted in **bold**. Predictive metrics: F1 (PTM, binding residue categorization) and AUROC (binding/catalytic site prediction).

Table 4 further extends the uncertainty quantification evaluation to NMV regression. As deterministic **UPLiFT** produces only point predictions, it can be evaluated using predictive metrics, but not distributional metrics such as CRPS and ECE. Hence, using a weight-space uncertainty baseline, we obtain predictive samples from **LoRA tuning** using Monte Carlo dropout (Gal & Ghahramani, 2016). Bayesian **UPLiFT** achieves lower CRPS and ECE than the MC dropout baseline, indicating improved distributional quality and calibration with a modest trade-off in predictive performance relative to its deterministic counterpart.

**Table 4:** Uncertainty quantification results for NMV. Point metrics use the predictive mean (20 samples for Bayesian **UPLiFT** and **LoRA tuning** MCD). Deterministic **UPLiFT** has no predictive distribution, so uncertainty metrics are omitted.

| Method | Predictive |  | Uncertainty |  |
| --- | --- | --- | --- | --- |
|  | MAE ↓ | Pearson ↑ | CRPS ↓ | ECE ↓ |
| Det. UPLiFT | **0.0387±0.0007** | **0.6738±0.0035** | — | — |
| Bayesian UPLiFT | 0.0399±0.0011 | 0.6527±0.0101 | **0.0410±0.0009** | **0.3935±0.0032** |
| LoRA tuning | 0.0391±0.0006 | 0.6721±0.0053 | 0.0433±0.0007 | 0.4861±0.0019 |

Overall, these results show that stochasticity over the activation parameters can enhance predictive calibration while preserving most of the performance of deterministic **UPLiFT**.

### 3.3 Activation-only Comparison with ct UNDER Frozen weights

To isolate the adaptation contributed by the randomly initialized weights, we freeze the pretrained PLM in its entirety and optimize only the activation parameters of the prediction head. Consequently, both methods (**UPLiFT** and **CT**) operate on the same fixed PLM representations and can adapt the downstream predictor only through their activation parameters.

**UPLiFT** consistently outperforms the **CT** parameterization across all four tasks (Table 9). In particular, **CT** produces negligible F1 scores at the fixed decision threshold. Even with retrospectively optimized thresholds, it achieves F1_max_ scores of 0.023, 0.122, 0.230, and 0.251 on PTM, binding-residue categorization, binding-site, and catalytic-site prediction, respectively, which are significantly below those of **UPLiFT** (0.175, 0.282, 0.382, and 0.512). This substantial gap is consistent with the behavior observed in the controlled toy example (Figure 1). Complete results across all metrics, together with additional experiments, are provided in Appendix E.

## 4 Learned representation across adaptations

Given the parity between LoRA and our approach in performance scale for the PTM multi-label dataset, one expects the learned residue representations, with reference to the pretrained model, to be similar for both approaches. Indeed, the UMAP visualization (McInnes et al., 2018) in Figure 3 illustrates broad similarity in the way both approaches organized the PTM-residue embedding. For example, the residues for Phosphorylation and O-linked Glycosylation are clustered together in both fine-tuned models, which is expected since both PTM groups have Serine (S) and Threonine (T) in common. Similar divergence from the pretrained representation is observed for multi-PTMs involving Acetylation. Specifically, note the concentrated region of Ace-associated multi-PTM residues in the pretrained model, and how both fine-tuned models form more distinctive regions. Such 2D projection-based similarity is further supported by the linear centered kernel alignment-based (Kornblith et al., 2019) comparison across the learned representation spaces (Figure 4, Appendix H.1).

**Figure 3:**
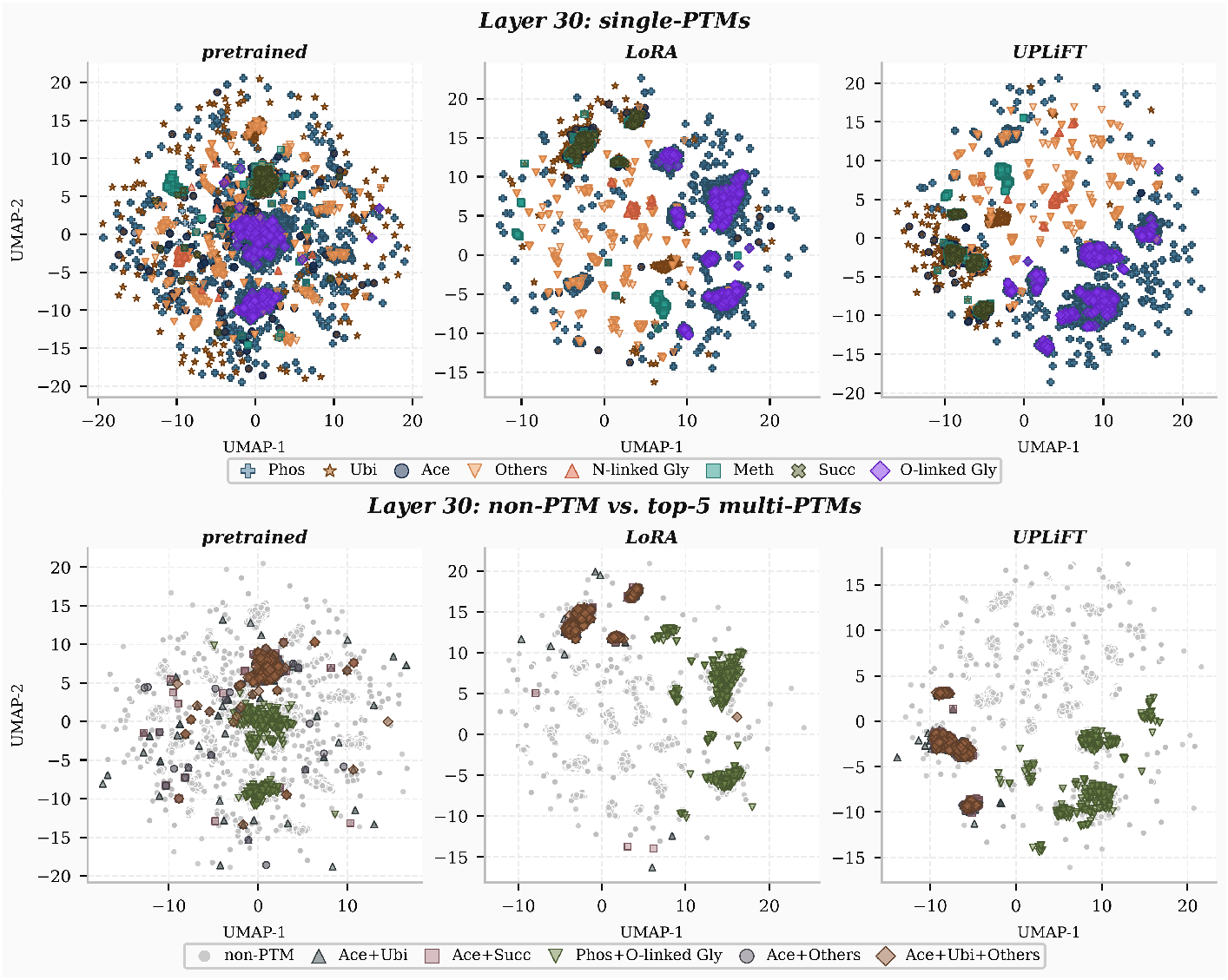
Residue representation from fine-tuned and pretrained models. The top panel shows UMAP residue embeddings (last layer) for sites with single positive PTM annotation. In the bottom panel, we have the top-5 multi-PTM sites (within the test-set) with non-PTM residues as background.

**Figure 4:**
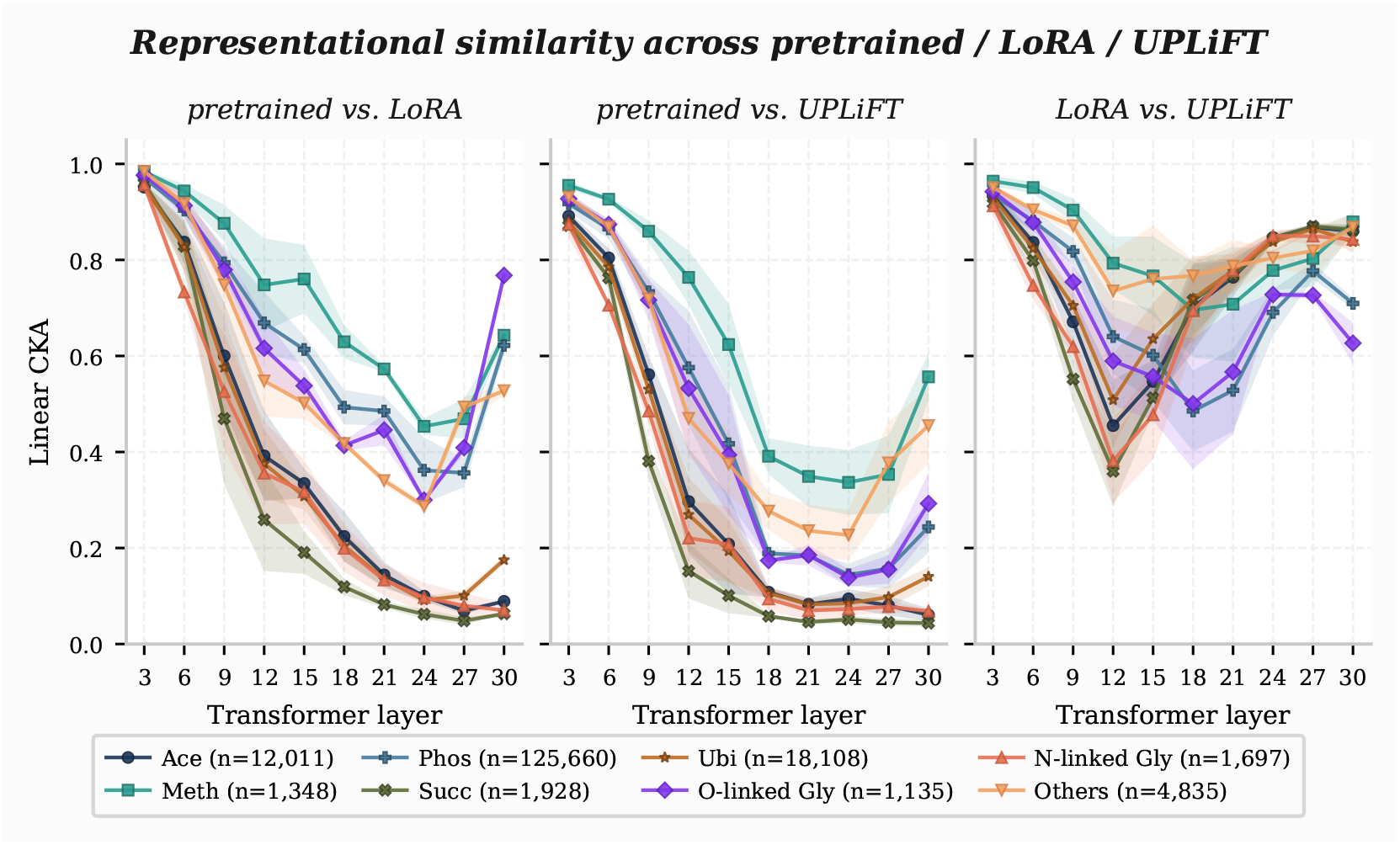
Trend of Linear CKA between representations from fine-tuned and pretrained models. The similarity (mean and standard deviation) is computed from fine-tuned models corresponding to 3 seeds reported in Table 2.

Since the most frequent PTM labels (excluding the rare groups) are associated with a limited set of amino acids, we wondered whether the fine-tuned models learned to organize the residues based on the amino acid identity. Our embedding separability analysis (Appendix H.2) revealed this trend for LoRA fine-tuned model, i.e., the silhouette coefficient (Euclidean distance) for AA identity-based clusters increased to 0.2783 ±0.0025 from 0.182 for the pretrained model (Table 13). In contrast, our proposed activation-only adaptation reshaped the embedding space with reduced AA identity-based separability (0.1473 ± 0.0217) while following LoRA fine-tuned representation space’s clusters based on physicochemical and PTM labels. This implies our approach did not drive the PLM to associate the samples with their PTM residue identity, which is not necessarily required for PTM classification. For instance, the hydroxyls of Serine and Threonine are key sites for Phosphorylation and O-linked Glycosylation, and both amino acids are in the same physicochemical group. Hence, fine-tuning with PTM multi-label classification is not intended to adapt the pretrained model to discriminate between these two residues, and UPLiFT representations showed this intuition.

## 5 Discussion and Limitations

In this work, we introduce UPLiFT, an activation-space-based parameter-efficient fine-tuning method with built-in uncertainty quantification. Across diverse protein prediction tasks, UPLiFT achieves competitive performance relative to weight-space baselines while requiring approximately 3 − 11× fewer trainable parameters. Bayesian UPLiFT further improves mean calibration and probabilistic metrics while largely retaining the predictive performance of deterministic UPLiFT. Our representation analysis for PTM classification task also reveals broad similarities between the residue embeddings learned by UPLiFT and LoRA. In particular, LoRA tuning strengthens AA-identity organization in the pretrained model during PTM fine-tuning, whereas UPLiFT achieves comparable PTM-label and physicochemical clustering without further emphasizing amino-acid separation. This suggests that UPLiFT exploits capabilities already present in the pretrained representation while concentrating its adaptation on the downstream task.

The limitations of our work can be summarized as follows. Although UPLiFT provides competitive performance relative to weight-space fine-tuning baselines, it generally performs below LoRA across both residue- and sequence-level tasks. On sequence-level tasks, UPLiFT also performs slightly below the Head tuning baseline. Although this trade-off is compensated by the substantially smaller number of trainable parameters, the remaining predictive gap is a limitation of the current formulation. A richer activation parameterization capable of expressing more complex nonlinearities may help reduce this gap. We further analyze how UPLiFT scales with the label-space size in Appendix I. While UPLiFT maintains strong performance as the number of labels increases, its gap from the weight-space baselines becomes larger. Finally, to isolate the benefits of activation-space fine-tuning and enable a controlled comparison across adaptation methods, we restricted our experiments to a fixed prediction architecture and PLM backbone. Investigating the effects of different architectures and PLM configurations remains an important direction for future work.

## Acknowledgments

This work was supported by the Advanced Research Projects Agency for Health (ARPA-H) Award 1AY1AX000053-01 and the National Science Foundation (NSF) Award IIS-2212419. Portions of this research were conducted with the advanced computing resources provided by Texas A&M High Performance Research Computing.

## A Related work

### Parameter efficient fine-tuning & uncertainty quantification

Pretrained foundation models provide a strong basis for downstream tasks in a specific domain, such as language (Yang et al., 2025; Touvron et al., 2023), vision (Bai et al., 2023), or proteins (Lin et al., 2023; Hayes et al., 2025; Candido et al., 2026; Elnaggar et al., 2022; 2023). Efficient adaptation to such tasks has been of paramount interest to the community in recent years. Lian et al. (2022) introduce an approach that adapts a pretrained model by learning feature-wise shift and scale parameters. LoRA (Hu et al., 2022) hypothesizes that updates to the pretrained model have a low intrinsic rank. Hence, while keeping the pretrained weights fixed, it formulates the updates as the product of two learnable low-rank matrices. Liu et al. (2024) decompose pretrained weights into magnitude and direction and apply low-rank adaptation to the directional component.

LoRA’s efficient formulation has also enabled uncertainty-aware fine-tuning for more reliable predictive models. Yang et al. (2024) introduce Laplace-LoRA, which applies a Laplace approximation to the posterior over the LoRA parameters. Wang et al. (2024) introduce a structured Bayesian framework to learn a posterior distribution over the low-rank matrices. Rahmati et al. (2026) further extend this formulation to model data-dependent uncertainty. Building upon this, Zhang et al. (2026b) provide a scalable Gaussian process-based framework for uncertainty-aware fine-tuning.

### Activation space adaptation

Hu et al. (2026) proposed curvature-tuning to smoothly alter each neuron’s decision boundary in a pretrained neural network by optimizing the curvature of the activation functions without changing the weights. Importantly, they performed this curvature optimization for the pretrained network while learning the weights in the task-specific prediction head with a fixed activation function. As we have demonstrated in Sections 2 and 3.3, this formulation can only steer neurons by scaling but lacks the affine-like control due to the frozen weights, thereby limiting its fine-tuning ability when the frozen weights are poorly suited for the prediction tasks. To address this issue, we propose a parameterized activation function that can adapt the neural network’s output more flexibly, even when the weights are frozen.

The utility of PLM’s activation space has gained more traction in protein engineering, as it allows a training-free approach to optimize protein sequence generation using a pretrained model. In particular, prior applications directly operating on the activation space of protein language models are aimed toward protein generation by steering either the raw hidden representation from PLM (Huang et al., 2025) or features learned through a sparse auto-encoder (SAE) (Simon & Zou, 2025). In contrast to this generative application, our work focuses on improving the discriminative power of a pretrained protein language model by tuning the proposed parameterized activation function without altering any weights in the prediction network–PLM with prediction head.

## B Details of datasets

### B.1 Sources

#### Post-translational modification (PTM)

We have used the train-validation-test splits of the multi-label dataset from Zhang et al. (2026a), deposited at https://zenodo.org/records/18389225. From dbPTM database (Chung et al., 2025), the authors curated PTM samples with 8 PTM labels: acetylation, methylation, N-linked glycosylation, O-linked glycosylation, phosphorylation, succinylation, ubiquitylation, and *“Others”*, which contains rest of the less-frequent PTM annotations. The splits were formed based on 40% sequence identity clustering via MMSeqs2 (Steinegger & Söding, 2017). Note each sample is 50 residue-long peptide with 8 binary multi-labels per site, where positive label indicates presence of each modification site.

#### Normal mode vector

Ni & Buehler (2026) developed a protein dataset of 12,924 samples with low-frequency normal modes to facilitate their generative approach for protein dynamics-targeted sequence design. We leverage their released dataset as residue-level regression task. Briefly, the normal mode vector for a sequence of N residues can be represented as ***v***_A_ = [*d*_1_, *d*_2_,*…, d*_*N*_] with 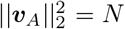, where each *d*_*i*_ is amplitude of displacement vector at *i*^th^ residue, extracted from normal mode analysis (NMA) (Hayward & de Groot, 2008) of relaxed protein structure. The detailed protocol, e.g., MD relaxation and NMA, can be found in Ni & Buehler (2026).

The rest of the prediction tasks are taken from the Magneton evaluation set (Calef et al., 2026), where further details can be found. Here we provide a brief description of these datasets.

#### Binding residue categorization

This residue-level multi-label classification task involves predicting each site’s binding partner: metal ions, nucleic acids, and small molecules.

#### Functional site prediction

This covers two residue-level binary classification tasks: ligand binding and catalytic sites in the sequence.

#### Binary and subcellular localization

These two sequence-level tasks are binary and multi-class classifications, respectively, for predicting the protein’s localizations. For binary localization, the goal is to classify whether the given protein is membrane-bound or soluble. The multi-class variant involves 10 subcellular classes.

#### Thermostability

The target of this sequence-level regression dataset is the protein’s melting temperature.

### B.2 Pre-processing

For the residue-level classification tasks, we followed the subsequence approach from Zhang et al. (2026a) for representing the samples as subsequences instead of providing full sequence to the model. Specifically, the subsequences are formed by padding the core region with 10 residues in both ends (20 in total) where core is selected with at least one residue site with positive label. The padding residues are taken from full sequence if sequence length permits, otherwise ‘X’ is used. The length of core varies across datasets: 50 (PTM, catalytic site prediction), 40 (binding site prediction), and 30 (binding residue categorization).

Since the normal mode vector data, accessible from supplemental information of Ni & Buehler (2026), does not have any fixed partitioning, we constructed train-validation-test split via MMseqs2 (Steinegger & Söding, 2017) with `min seq id=0.4`. Unlike other residue-level tasks, we utilized full sequence for this property since the ground truth vectors were normalized with the sequence length.

We excluded any sequence longer than 1024 residues in all sequence-level tasks of Table 2 except for thermostability, where maximum sequence length cutoff of 1000 was used.

## C Evaluation metrics

### Performance metrics

Depending on the task, we evaluate predictive performance using several metrics that reflect different criteria. For classification tasks, F1 measures how well the model identifies positive labels while maintaining high precision and recall:

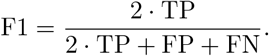

AUROC measures the model’s ability to rank positive samples above negative samples, while AUPRC evaluates the trade-off between precision and recall across thresholds:

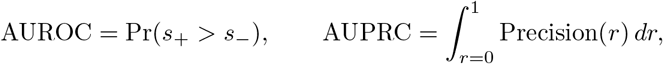

where *r* denotes recall and *s*_+_ and *s*_−_ denote the scores assigned to positive and negative samples, respectively. The Matthews correlation coefficient (MCC) further measures the correlation between binary predictions and true labels, rewarding correct predictions of both positive and negative samples while penalizing both types of error:

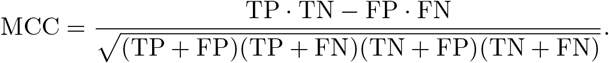

For classification tasks, we further evaluate accuracy.

For regression tasks, predictive performance is measured by Pearson correlation, mean absolute error (MAE), and root mean squared error (RMSE):

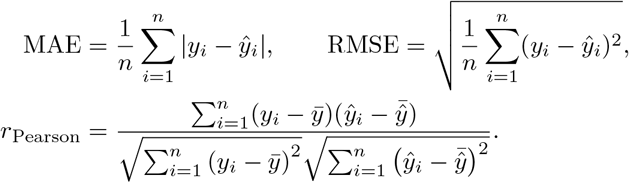

### Uncertainty metrics

We evaluate uncertainty quantification mainly by measuring how well calibrated the predictions are and how overconfident the model is in its predictions. For classification tasks, negative log-likelihood (NLL) penalizes incorrect predictions made with high confidence:

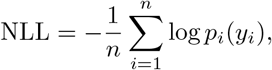

where *p*_i_(*y*_i_) denotes the probability assigned to the true label. The Brier score measures the squared difference between predicted probabilities and observed labels:

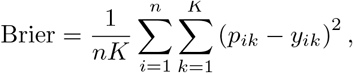

where *K* is the number of labels. Lower values indicate more accurate and better-calibrated probabilistic predictions. Expected calibration error (ECE), on the other hand, measures how closely the model’s prediction accuracy aligns with its confidence by binning the predictions and taking the weighted average of the difference between the two in each bin:

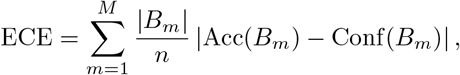

where

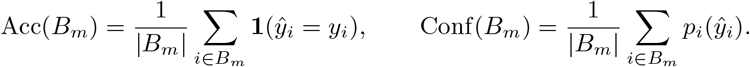

For regression tasks, the continuous ranked probability score (CRPS) evaluates the predictive distribution by comparing its cumulative distribution function with the observed target:

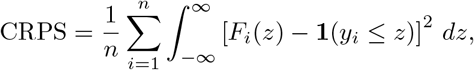

where *F*_*i*_ denotes the predictive cumulative distribution function for sample *i*. Lower CRPS values indicate better probabilistic predictions.

## D Detailed results for Table 2

Tables 5 and 6 show detailed performance metrics for the classification tasks in Table 2. Table 7 does the same

**Table 5:** Detailed results on residue-level classification tasks. The metrics in PTM and binding residue categorization tasks are computed as macro average.

| Task | Method | F1 (↑) | MCC (↑) | AUROC (↑) | AUPRC (↑) |
| --- | --- | --- | --- | --- | --- |
| PTM | LoRA tuning | **0.4089**±0.0037 | **0.4189**±0.0019 | **0.9807**±0.0005 | **0.3812**±0.0016 |
| | Head tuning | 0.3124±0.0020 | 0.3106±0.0025 | 0.9766±0.0005 | 0.2780±0.0026 |
| | UPLiFT | 0.4080±0.0026 | 0.4149±0.0019 | 0.9796±0.0005 | 0.3787±0.0030 |
| Binding Residue Categorization | LoRA tuning | **0.3327**±0.0061 | 0.3247±0.0029 | 0.8026±0.0084 | 0.3323±0.0119 |
| | Head tuning | 0.3116±0.0067 | **0.3297**±0.0013 | **0.8038**±0.0055 | **0.3445**±0.0054 |
| | UPLiFT | 0.3158±0.0074 | 0.3010±0.0020 | 0.7674±0.0037 | 0.2937±0.0031 |
| Binding Site Binary | LoRA tuning | **0.5522**±0.0004 | **0.5470**±0.0021 | **0.8990**±0.0011 | **0.6368**±0.0016 |
| | Head tuning | 0.3730±0.0147 | 0.3840±0.0066 | 0.8630±0.0016 | 0.4667±0.0028 |
| | UPLiFT | 0.4916±0.0040 | 0.4814±0.0073 | 0.8772±0.0013 | 0.5536±0.0091 |
| Catalytic Site Binary | LoRA tuning | **0.6563**±0.0070 | **0.6522**±0.0069 | **0.9371**±0.0012 | **0.7063**±0.0050 |
| | Head tuning | 0.5115±0.0111 | 0.5257±0.0079 | 0.9010±0.0047 | 0.5578±0.0109 |
| | UPLiFT | 0.5799±0.0178 | 0.5737±0.0149 | 0.9194±0.0014 | 0.6086±0.0136 |

**Table 6:**
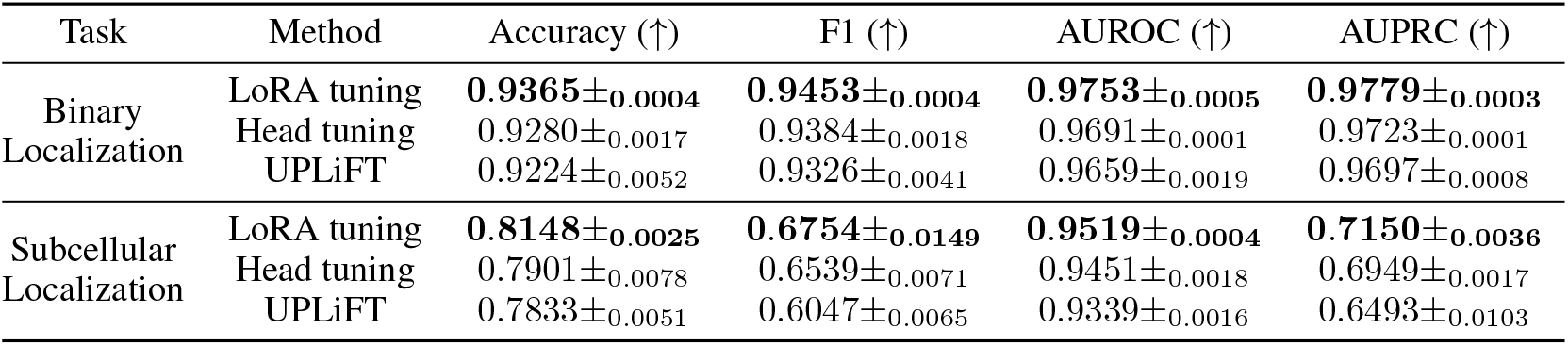
Detailed results on sequence-level classification tasks. The metrics (except for accuracy) in Subcellular localization task are computed as (OvR) macro average.

**Table 7:** Detailed results on regression tasks.

| Task | Method | Pearson ↑ | MAE ↓ | RMSE ↓ |
| --- | --- | --- | --- | --- |
| Normal Mode Vector | LoRA tuning | 0.6720±0.0053 | 0.0391±0.0006 | 0.0574±0.0005 |
| | Head tuning | 0.6353±0.0031 | 0.0420±0.0015 | 0.0604±0.0012 |
| | UPLiFT | **0.6738**±0.0035 | **0.0387**±0.0007 | **0.0573**±0.0005 |
| Thermostability | LoRA tuning | 0.6852±0.0074 | 0.5908±0.0092 | 0.7510±0.0099 |
| | Head tuning | **0.6894**±0.0063 | **0.5892**±0.0037 | **0.7461**±0.0016 |
| | UPLiFT | 0.6586±0.0078 | 0.6232±0.0041 | 0.7847±0.0056 |

## E Comparison with ct (Hu et al., 2026)

In this section, we compared **UPLiFT** and **CT** (Hu et al., 2026) in the activation-only fine-tuning approach for 4 residue-level classification datasets. For **CT**, we initialized PLM’s *ϕ*_CT_ with *t* = 0.64, *c* = 1, and prediction head’s *ϕ*_CT_ with *t* = *c* = 0.5. In **UPLiFT**, the PLM and prediction head’s nonlinearities are initialized with 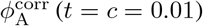 and *ϕ*_A_ (*t* = 0.5, *c* = 3), respectively^1^. These particular initializations were made to preserve the pretrained model’s activation function response at the beginning of the activation-only approach.

As we see from Table 8, fine-tuning via our **UPLiFT** led to better (or comparable) performance on the majority of the metrics across four datasets. For the multi-label classification tasks (PTM and binding residue categorization), *ϕ*_CT_ significantly falls behind **UPLiFT**. We hypothesized that the predictive performance of the **CT** was due to activation parameters for the pretrained weights in the PLM. This is supported by the comparison results with the frozen PLM (Table 9), as discussed in Section 3.3.

**Table 8:** Adaptation results for activation-only (PLM+prediction head) fine-tuning. All performance metrics are aggregated over three seeds. For PTM and binding residue categorization datasets, we report the macro-averaged scores. In each metric, the best method without overlap in mean ± standard deviations is highlighted.

| Task | Methods | AUROC (↑) | AUPRC (↑) | F1 (↑) | MCC (↑) | F1<sub>max</sub> (↑) |
| --- | --- | --- | --- | --- | --- | --- |
| PTM | det. UPLIFT | **0.9796±0.0005** | **0.3787±0.0030** | **0.4080±0.0026** | **0.4149±0.0019** | **0.4097±0.0024** |
| | CT | 0.9718±0.0041 | 0.3707±0.0047 | 0.3595±0.0051 | 0.3805±0.0040 | 0.3754±0.0033 |
| Binding Residue Categorization | det. UPLIFT | **0.7674±0.0037** | **0.2937±0.0031** | **0.3158±0.0074** | **0.3010±0.0020** | **0.3592±0.0074** |
| | CT | 0.7115±0.0293 | 0.2151±0.0342 | 0.2253±0.0127 | 0.2130±0.0246 | 0.2709±0.0321 |
| Binding Site Binary | det. UPLIFT | **0.8772±0.0013** | 0.5536±0.0091 | **0.4916±0.0040** | 0.4814±0.0073 | **0.5255±0.0050** |
| | CT | 0.8464±0.0099 | 0.5542±0.0140 | 0.4822±0.0086 | **0.5041±0.0069** | 0.5002±0.0121 |
| Catalytic Site Binary | det. UPLIFT | **0.9194±0.0014** | 0.6086±0.0136 | 0.5799±0.0178 | 0.5737±0.0149 | **0.5925±0.0124** |
| | CT | 0.8808±0.0051 | 0.5980±0.0125 | 0.5748±0.0118 | 0.5816±0.0137 | 0.5762±0.0120 |

**Table 9:** Adaptation results for activation-only (prediction head) fine-tuning. Unlike Table 8, we have optimized the activation parameters of only the prediction head while keeping the PLM frozen to pretrained configurations.

| Task | Methods | AUROC (↑) | AUPRC (↑) | F1 (↑) | MCC (↑) | F1<sub>max</sub> (↑) |
| --- | --- | --- | --- | --- | --- | --- |
| PTM | det. UPLIFT | **0.8346±0.0177** | **0.1334±0.0038** | **0.1524±0.0042** | **0.1475±0.0042** | **0.1754±0.0041** |
| | CT | 0.5106±0.0210 | 0.0123±0.0037 | 0.0000±0.0000 | -0.0001±0.0001 | 0.0225±0.0029 |
| Binding Residue Categorization | det. UPLIFT | **0.7336±0.0064** | **0.2209±0.0098** | **0.1096±0.0269** | **0.1632±0.0258** | **0.2822±0.0081** |
| | CT | 0.5173±0.0386 | 0.0760±0.0104 | 0.0339±0.0549 | 0.0096±0.0130 | 0.1216±0.0043 |
| Binding Site Binary | det. UPLIFT | **0.8039±0.0042** | **0.3410±0.0065** | **0.2387±0.0073** | **0.2661±0.0053** | **0.3821±0.0042** |
| | CT | 0.6353±0.0311 | 0.1613±0.0253 | 0.0000±0.0000 | 0.0000±0.0000 | 0.2300±0.0283 |
| Catalytic Site Binary | det. UPLIFT | **0.8950±0.0014** | **0.5116±0.0102** | **0.4457±0.0109** | **0.4716±0.0077** | **0.5117±0.0056** |
| | CT | 0.7067±0.0578 | 0.1828±0.0798 | 0.0000±0.0000 | 0.0000±0.0000 | 0.2514±0.0687 |

## F Ablation experiments for the correction term in 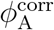

To investigate the importance of the non-trainable correction term in 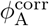, we performed the activation-only adaptation by replacing 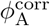with *ϕ*_A_, while keeping other training hyper parameters same as in the experiment for Table 2. The large performance drops in this setting (“wo correction (*t*=*c*=0.01)” in Table 10), compared to deterministic **UPLiFT**, highlight the need for preserving the activation response of the pretrained network at initialization of learnable activation parameters.

**Table 10:** Ablation of the correction term. The values for *c, t* denote the initialization of the activation parameters in PLM.

| Task | Metrics (↑) | det. UPLiFT<br>($t=c=0.01$) | wo correction<br>($t=c=0.01$) | wo correction<br>($t=0.5, c=1.0$) |
| --- | --- | --- | --- | --- |
| PTM | F1 | **0.4080**±0.0026 | 0.3434±0.0068 | 0.3976±0.0044 |
| | MCC | **0.4149**±0.0019 | 0.3594±0.0069 | 0.4036±0.0040 |
| | AUROC | **0.9796**±0.0005 | 0.9731±0.0017 | 0.9784±0.0003 |
| | AUPRC | **0.3787**±0.0030 | 0.3123±0.0122 | 0.3651±0.0011 |
| Binary<br>Localization | Accuracy | **0.9224**±0.0052 | 0.8069±0.0045 | 0.8598±0.0074 |
| | F1 | **0.9326**±0.0041 | 0.8430±0.0045 | 0.8800±0.0065 |
| | AUROC | **0.9659**±0.0019 | 0.8507±0.0009 | 0.9091±0.0032 |
| | AUPRC | **0.9697**±0.0008 | 0.8505±0.0029 | 0.9097±0.0031 |

**Table 11:**
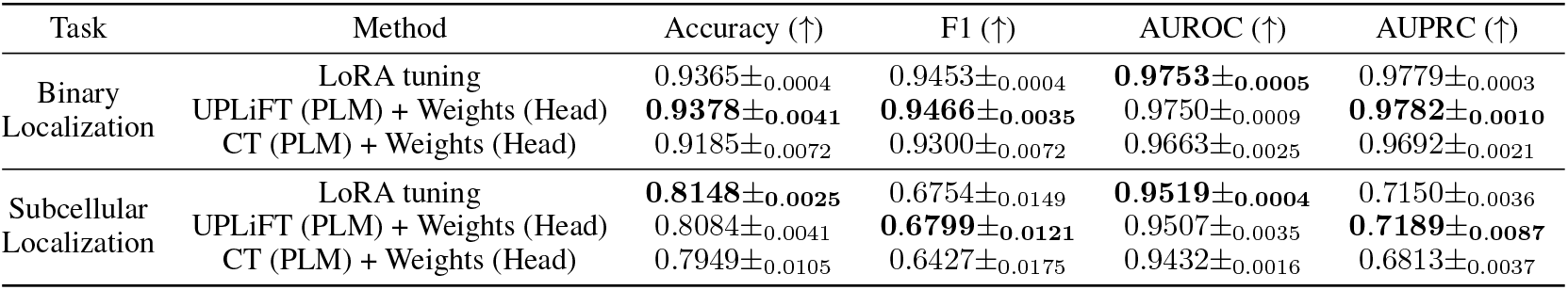
Detailed results for fine-tuning with activation in PLM and weights in prediction head on sequence-level classification tasks. The metrics (except for accuracy) in Subcellular localization task are computed as (OvR) macro average. The best mean performance metrics are highlighted in **bold**.

**Table 12:** Detailed results for fine-tuning with activation in PLM and weights in prediction head on Thermostability task (sequence-level regression).

| Task | Method | Pearson ↑ | MAE ↓ | RMSE ↓ |
| --- | --- | --- | --- | --- |
| Thermostability | LoRA tuning | **0.6852**±0.0074 | **0.5908**±0.0092 | **0.7510**±0.0099 |
| | UPLiFT (PLM) + Weights (Head) | 0.6788±0.0033 | 0.5958±0.0046 | 0.7554±0.0053 |
| | CT (PLM) + Weights (Head) | 0.6776±0.0040 | 0.5998±0.0048 | 0.7651±0.0062 |

Without the correction term, we have empirically observed that the initialization *t* = 0.5 and *c* = 1.0 −3.0 in the PLM–similar to the values used in the prediction head–leads to better predictive performance, but it still falls short of what the correction term achieves. Particularly, fine-tuning from this specific initialization (“wo correction (*t*=0.5, *c*=1.0)”) brings performance on the PTM dataset very close to deterministic **UPLiFT** with the correction term, while the gap is still moderate in the binary localization dataset. This discrepancy likely reflects the fact that, for the latter dataset, the pretrained model already offers strong discriminative power without further fine-tuning–as illustrated by **Head tuning**’s performance (Table 2)–so the correction term plays a more significant role here by allowing activation-only adaptation to effectively leverage the pretrained PLM representation.

## G Learning prediction head weights in sequence-level tasks

In contrast to the activation-only adaptation results in the main text (Table 2), this section reports prediction performance for fine-tuning with **UPLiFT**/**CT** in the PLM while learning the prediction-head weights for sequence-level tasks. These two activation approaches have the same fixed activation ReLU in the prediction head as in the **LoRA tuning**. As we see from Table 11, our proposed activation is competitive with LoRA while outperforming **CT** in both classification tasks. Table 12 extends this observation to the Thermostability regression task where **UPLiFT** consistenly outperforms **CT** while offering competitive performance with LoRA. This result further demonstrates not only the importance of prediction head weights for the global property in sequence-level tasks, but also **UPLiFT**’s improved adaptation capacity over **CT** under pre-trained PLM weights.

## Is UPLiFT adapting the PLM in the same manner as LORA?

### H.1 Representation similarity

Complementary to the 2D-UMAP visualization of fine-tuned representation for PTM dataset (Figure 3), we further quantified the change in residue representation space across multiple layers {3, 6,*…*, 30}of PLM via linear centered kernel alignment (CKA,Kornblith et al. (2019)). Specifically, we compared the embeddings for each group of PTM residues, i.e., sites annotated with the positive label for the corresponding PTM in the dataset’s test split.

As seen from Figure 4, both LoRA and our approach gradually diverge away from pretrained representation in mostly similar fashion except for the relative differences across the PTM labels. For instance, the representation spaces for Phosphorylation and O-linked Glycosylation moved further away in our activation-only approach, whereas LoRA pulled them toward a high-similarity (with respect to pretrained) region in the last layer. Nonetheless, both fine-tuned representations for these two groups are still highly similar (linear CKA above 0.6 in final layers), albeit with lower similarity than that of remaining PTM groups.

Another distinct characteristic of our approach is the diversity of the fine-tuned representations for different PTM groups, indicated by the higher standard deviations of representational similarity across 3 seeds of fine-tuned models. Specifically, the LoRA fine-tuned representations for all three seeds converged to consistently similar region for all PTM groups, whereas our activation-only approach showed PTM-label sensitivity. For instance, across multiple seeds the samples from “Others”, i.e., combination of all rare PTM groups, were adapted in different degree of similarity with respect to the pretrained model. Since all rare PTM samples are grouped into this same label, it is more intuitive that the fine-tuned models are diverse in encoding such group, hinting at our approach’s potential in learning informative representation uncertainty.

### H.2 Rearrangement of PLM REPRESENTATIONS

To gain insights into the mechanism that the fine-tuning approaches followed to update the residue representation, we analyzed the cluster separability of the positive samples from the 7 most-frequent single-PTM residues. Specifically, we computed the silhouette coefficient (Rousseeuw, 1987) for three grouping criteria: amino acid identity (`K,L,N,R,S,T,Y`), physicochemical properties (aromatic, hydrophobic, polar, positively charged), and the PTM labels.

For the LoRA fine-tuned model, the separability quality for these three criteria increased from that of pretrained representation (Table 13). On the other hand, our activation-only approach matches the clustering quality of LoRA for both physicochemical and PTM grouping, while it leads to reduced separation for amino acid identities. This is particularly interesting since it shows that it is not necessary to associate the samples with their PTM residue identity to achieve comparable predictive performance. Specifically, these PTMs share similar amino acid preference confined to the narrow set of 7 amino acids. Since the positive samples in dataset are dominated by the high-frequency PTM groups, it could be convenient for the fine-tuning approaches to leverage such characteristics of the dataset to adapt the pretrained representation. The discrepancy in AA identity criterion between our approach and LoRA illustrates that the activation-only adaptation rearranged the residue embedding space of the pretrained model, necessitated by the task-specific loss with reduced sensitivity to the imbalanced composition of the dataset.

## I Analysis of Label-Space scaling

We further examine how activation-only adaptation scales as the number of output labels increases. This setting can particularly be challenging for **UPLiFT** because the task-specific output projection remains randomly initialized and frozen. Unlike **Head tuning** and **LoRA tuning, UPLiFT** cannot update the output weights for each label; instead, its shared activation parameters must simultaneously adapt the representations to support an increasing number of ouptut labels.

Table 14 compares the top-4 and top-50 settings, i.e., classification for 4 and 50 most frequent labels, for GO-CC (Gene Ontology annotations for Cellular Component) and EC (Enzyme Commission) (Gligorijević et al., 2021). **UPLiFT** remains competitive in the top-4 setting, but its gap from the strongest weight-space baseline increases for top-50. For GO-CC, the AUPRC gap increases from 0.024 to 0.091, while the F1 gap increases from 0.019 to 0.051. Similarly, for EC, the AUPRC gap increases from approximately 0.007 to 0.041, and the F1 gap increases from 0.017 to 0.044. Nevertheless, **UPLiFT** retains strong absolute performance on top-50 EC, achieving an AUROC of 0.979, an AUPRC of 0.911, and an F1 score of 0.861.

**Table 13:** Silhouette coefficient (↑ : higher degree of separation across groups) for 7 most-frequent single-PTM samples. Residue embeddings from last layer of each model are extracted for around 50000 stratified samples, with minimum of 1000 samples per group. For each seed, the scores are averaged over 5 repeats of sub-sampling (standard deviation was ≈0.001), and the results are reported for fine-tuned models from 3 seeds. The cases with substantial difference between LoRA and our approach are highlighted in **bold**.

| Grouping | Cosine |  |  | Euclidean |  |  |
| --- | --- | --- | --- | --- | --- | --- |
|  | LoRA | Ours | Pretrained | LoRA | Ours | Pretrained |
| AA identity | **0.4630 ± 0.0066** | 0.2327 ± 0.0395 | 0.312 | **0.2783 ± 0.0025** | 0.1473 ± 0.0217 | 0.182 |
| Physicochemical | 0.3400 ± 0.0204 | 0.3280 ± 0.0731 | 0.231 | 0.2040 ± 0.0125 | 0.2080 ± 0.0466 | 0.136 |
| PTM label | 0.0783 ± 0.0291 | 0.0717 ± 0.0280 | 0.042 | 0.0433 ± 0.0147 | 0.0403 ± 0.0132 | 0.024 |

**Table 14:** Effect of label-space size. Top-4 numbers are mean ±std over three runs. Top-50 is a single run. Best result in each column is in **bold**.

| Task | Method | Top-4 |  |  | Top-50 |  |  |
| --- | --- | --- | --- | --- | --- | --- | --- |
|  |  | AUROC | AUPRC | F1 | AUROC | AUPRC | F1 |
| GO-CC | Head tuning | 0.808±0.003 | 0.812±0.003 | 0.687±0.014 | **0.864** | **0.616** | 0.469 |
|  | LoRA tuning | **0.809±0.003** | **0.815±0.003** | **0.702±0.005** | 0.852 | 0.592 | **0.471** |
| | UPLiFT | 0.789±0.008 | 0.791±0.006 | 0.683±0.003 | 0.835 | 0.525 | 0.420 |
| EC | Head tuning | **0.9996±0.0001** | **0.9989±0.0002** | **0.985±0.005** | **0.994** | **0.952** | 0.903 |
| | LoRA tuning | 0.9985±0.0012 | 0.995±0.003 | 0.973±0.008 | 0.992 | 0.948 | **0.905** |
| | UPLiFT | 0.997±0.002 | 0.992±0.006 | 0.968±0.004 | 0.979 | 0.911 | 0.861 |

These results highlight scaling to larger label spaces as an important direction for further study. **UPLiFT** remains effective in the top-50 setting, while the observed differences suggest opportunities for further improvement as the number of labels increases. Since the top-50 results correspond to a single run, they provide preliminary evidence of this scaling behavior.

## J Analysis on UPLiFT derivatives

### Lemma J.1

(Controlled input sensitivity and local curvature). *For fixed t* ∈ (0, 1) *and c >* 0, *define*

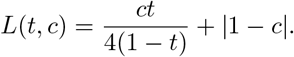

*Then ϕ*_A_(·; *t, c*) *is globally L*(*t, c*)*-Lipschitz:]*

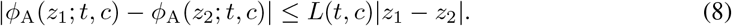

*Moreover, on every open interval where u*(*z*; *t, c*) *>* 0,

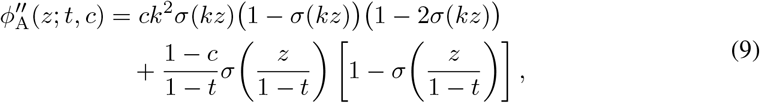

*where k* = *t/*(1 − *t*), *and therefore*

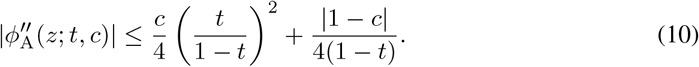

*Proof*. On an interval where *u >* 0, *ϕ*_A_ = *u*. Differentiating *u* gives

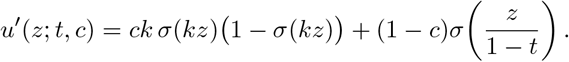

Using 0 ≤*σ*(*v*)(1 − *σ*(*v*)) ≤1*/*4 and 0 ≤ *σ*(*v*) ≤1 yields |*u*^′^(*z*; *t, c*) |≤*L*(*t, c*) uniformly in *z*. Since the ReLU is 1-Lipschitz, its composition with *u* is also *L*(*t, c*)-Lipschitz, proving Equation (8).

Differentiating once more gives Equation (9). Applying |1 −2*σ*(*v*) | ≤1 and *σ*(*v*)(1 −*σ*(*v*)) ≤1*/*4 gives Equation (10).

Definition 2.2 characterizes the additional adaptation afforded by UPLiFT through its rectified-affine tail, whereas Definition J.1 shows that this flexibility does not cause the input gradient to grow with | *z* |. At the same time, Equation (9) remains nonzero in general, allowing UPLiFT to modify local curvature while the remaining model parameters are fixed.

### Corollary J.2

(Sensitivity of the corrected activation). *Let*

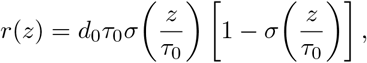

*where d*_0_, *τ*_0_ *>* 0, *such that*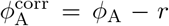. *For fixed t* ∈ (0, 1) *and* 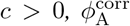*is globally Lipschitz with constant*

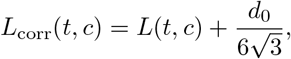

*where*

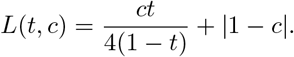

*Moreover, away from the switching points u*(*z*; *t, c*) = 0,

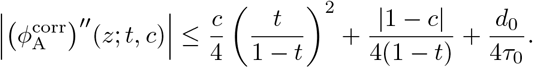

*Proof*. Let *p*(*z*) = *σ*(*z/τ*_0_). Differentiating the correction gives

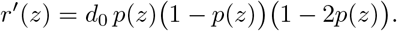

For *p* ∈ [0, 1],

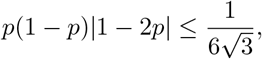

and hence

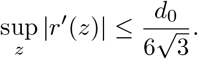

The Lipschitz result follows from 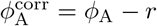 and Definition J.1.

Differentiating once more yields

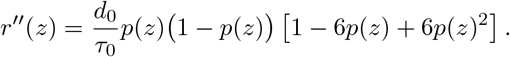

Since

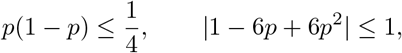

we have

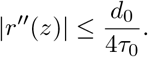

Combining this result with the curvature bound in Definition J.1 proves the claim.

## K Approximating GELU With 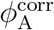

By setting *t* = *c* = 0 in Equation (4), we opted to find the parameters *d*_0_, *τ*_0_ to introduce the negative response region near *z* = 0. Accordingly, we applied global optimization (via `scipy.optimize.differential evolution` (Virtanen et al., 2020)), followed by local refinement (via `scipy.optimize.minimize)`, to minimize approximation error (Equation (11)) between GELU and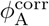.

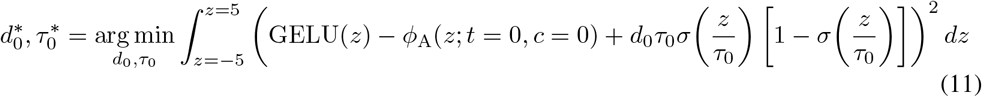

## L Training details

All models are optimized using Adam (Kingma & Ba, 2015) with batch sizes of 256 (residue-level classification), 32 (sequence-level classification), and 64 (regression). For all binary classification tasks, we use binary cross-entropy as task loss, except for PTM, where we use a hybrid loss combining Dice and focal losses with a probability magnitude penalty, following Zhang et al. (2026a). For subcellular localization (multiclass classification), the task loss is cross-entropy with a label-smoothing factor of 0.1. Both regression tasks use MSE as the task objective, with an additional vector-norm penalty for normal-mode vector prediction.

Hyperparameters were selected separately for each task based on validation performance. We use F1 for multi-label classification, AUROC for binary and multi-class classification, and Pearson correlation for regression. For the **LoRA tuning** and **Head tuning** baselines, we follow the default settings (e.g., LoRA rank, prediction head architecture) used by Zhang et al. (2026a), with task-specific learning rates reported below.

### Deterministic experiments

For LoRA tuning, we use a learning rate of 2 ×10^−5^ for all tasks except binding-residue categorization, where we use 10^−4^. For Head tuning, the learning rate is 10^−5^ for PTM and localization tasks and 2 ×10^−4^ for binding-residue, binding-site, and catalytic-site prediction.

For deterministic UPLiFT, we use lr = 5 ×10^−3^ across classification tasks, except for binding-residue categorization, where we use 10^−2^. The trainable parameters (*a, b, c*) of UPLiFT are initialized in such manner that we have (*c, t*) = (0.01, 0.01) in the PLM and (*c, t*) = (3.0, 0.5) in the prediction head.

For regression tasks, the learning rates for *a, b* (corresponding to *t*) and *c* are 10^−3^ and 2.5 ×10^−3^, respectively. Furthermore, we initialize the UPLiFT parameter *t* to 0.1 for both PLM and the prediction head. For the scaler parameter *c* in PLM, we have *c* = exp(−4) for NMV and thermostability regression. Corresponding initialization values for *c* in the prediction head are exp(1) and exp(3).

We train for 100 epochs for all datasets, except for normal-mode vector regression, where we let it run for 300 epochs.

#### Bayesian experiments

For Bayesian UPLiFT, activation parameters are sampled during training, while checkpoint selection is performed using the distribution mean. The concentration scale is fixed at *κ* = 2 across all tasks. For classification tasks, we use a learning rate of 5 ×10^−3^. The normal-mode vector experiment uses the same setups as its deterministic counterpart. Furthermore, all initializations of UPLiFT parameters are the same for both deterministic and Bayesian variants.

We use a Beta(5, 5) prior for PTM and catalytic-site prediction, a Beta(10, 10) prior for binding-site, Beta(1, 9) normal-mode vector prediction, and a Beta(1, 10) prior for binding-residue categorization. We set *λ* = 0.75 for PTM, NMV, and catalytic-site prediction and *λ* = 0.95 for the remaining tasks. The KL warmup lasts 100 epochs for PTM and catalytic-site prediction, 50 epochs for binding-residue and binding-site prediction, and 20 epochs for normal-mode vector prediction. We use *γ* = 10^−6^ for binding-residue and catalytic-site prediction, 5 × 10^−6^ for PTM, 3 ×10^−6^ for binding-site prediction, and 5 ×10^−4^ for normal-mode vector prediction. Finally, training is performed for 200, 300, 100, 500, and 1500 epochs for PTM, binding-residue, catalytic-site, binding-site, and normal-mode vector prediction, respectively.

#### Computational resource

All experiments were conducted using a single NVIDIA H200 GPU with 141GB.

## Footnotes

1 In our implementation, we parameterize *c* using an exponential function to ensure positivity.

